# SplitAligner: A Gene-Species Tree Reconciliation Framework Using Split-Based Branch Mapping

**DOI:** 10.64898/2026.02.24.707838

**Authors:** Jiaqi Wu

## Abstract

Phylogenomic analyses are increasingly focused on branch-specific questions within a fixed species tree. However, two pervasive challenges in real datasets—missing taxa and gene-tree/species-tree discordance—complicate the comparability of branches across loci. Here, we introduce SplitAligner, a split-based framework that defines branch identity on a fixed species-tree backbone and evaluates it gene by gene under varying taxon coverage. For each species-tree branch, SplitAligner projects its split onto the gene-specific taxon set to determine whether the branch is evaluable or structurally missing due to a degenerate projected split. Under fixed-topology gene trees, this projection reveals branch fusion, where multiple species-tree branches collapse into an indistinguishable fusion group on the observed taxa. SplitAligner reports such cases as composite fused-branch identities and aggregates branch lengths accordingly. Under free-topology gene trees, SplitAligner further identifies topology-induced missingness (NA_topo), where a branch is decisive under the projected-split criterion but its projected split is absent from the gene tree, separating discordance-driven absence from coverage-driven missingness. These operations produce standardized gene × branch tables and a branch-wise concordance score (Support), defined as the fraction of decisive genes whose free-topology trees recover each projected split. Applying SplitAligner to 2,275 single-copy genes from a dataset of 302 mammals reveals heterogeneous concordance across the mammalian phylogeny and highlights internodes with elevated discordance-associated missingness. The resulting branch coordinate system provides a general framework for branch-based estimates of evolutionary rates, selective constraints, and other branch-wise summaries across thousands of loci/genes.

## 1. Introduction

Phylogenomic studies increasingly ask branch-specific questions on a fixed species tree, yet two pervasive properties of real datasets—gene-tree-discordance and missing taxa—complicate the comparability of branches across loci. Discordance is expected under the multispecies coalescent and is especially pronounced across short internodes, where incomplete lineage sorting generates alternative gene-tree resolutions even without error (*1, 2*). Missing data are similarly widespread in large genomic resources and can affect phylogenomic inference and downstream summaries (*3*). Together, discordance and missing taxa make “branch support” ambiguous unless branch identity and the causes of missing mappings are explicitly defined under gene-specific taxon sets.

A common strategy for enabling branch-wise comparisons across genes is to constrain gene trees to a reference species-tree topology and estimate gene-wise branch lengths on a fixed backbone (*4*). This provides a convenient coordinate system, but it can also obscure discordance-driven signals by construction and may introduce systematic biases when locus histories deviate from the fixed topology (*5*). Conversely, free-topology gene trees preserve locus-specific histories, but mapping their branches onto a shared species-tree coordinate system becomes non-trivial when missing taxa cause projected branches to collapse or become ambiguous. These limitations motivate an explicit branch-identity infrastructure that remains well-defined under both missing taxa and gene-tree/species-tree discordance.

Several methods summarize concordance across genes, including concordance factors in IQ-TREE2 (gCF/sCF) (*6*) and local branch support measures from coalescent-based species-tree inference (e.g., ASTRAL local posterior probabilities) (*7*). The most closely related split-based tool is PhyParts (*8*), which maps gene-tree bipartitions to a reference species tree to summarize concordant and conflicting splits; it also notes that missing taxa can lead to ambiguous multi-mapping of bipartitions, typically treated as ambiguity or down-weighted support. Here we introduce SplitAligner, which reframes multi-mapping as a projection-induced branch fusion phenomenon and provides a branch-coordinate infrastructure: projected splits define branch identity on gene-specific taxon sets, missing outcomes are decomposed into NA_struct / NA_fuse / NA_topo with an explicit accounting identity, and mapping results are exported as standardized gene×branch tables together with an interpretable branch-wise concordance score (Support) for downstream branch-wise analyses.

We make three contributions.

1. We define a **split-based branch coordinate system** that preserves branch identity under gene-specific taxon sets, including explicit representation of fused branches under missing taxa.
2. We provide a **missingness decomposition and accounting framework** (NA_struct / NA_fuse / NA_topo) that separates taxon-coverage effects from discordance-driven absence and yields internal consistency checks for branch-wise mapping.
3. We introduce a branch-wise **concordance annotation (Support)** that summarizes how frequently each species-tree internode is recovered across free-topology gene trees among decisive loci, enabling direct visualization and quantification of discordance hotspots on the species-tree backbone.

## 2. Overview and Algorithm

### 2.1 Overview of the split-based branch coordinate system

SplitAligner defines branch identity on a fixed species-tree backbone using splits (bipartitions) and evaluates that identity gene by gene under varying taxon coverage. Let *S* denote the species tree with taxon set *T*, and let *G_g_* denote the gene tree for gene *g* with observed taxon set *T_g_* ⊆ *T*. Each internal branch *b* of *S* induces a species-tree split σ(*b*) = (*A_b_* | *B_b_*). To compare this branch across genes with missing taxa, SplitAligner constructs the **projected split** on the gene-specific taxon set,

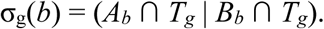

A projected split is **degenerate** if it collapses to a trivial partition (e.g., one side becomes empty) due to insufficient taxon coverage. In that case, the branch is not evaluable for gene *g*, producing **structural missingness**. Otherwise, gene *g* is **decisive** for branch *b*, and SplitAligner determines whether the projected split σ_g_(*b*) is present among the bipartitions induced by the gene tree (i.e., σ_g_(*b*) ∈ Σ(*G_g_*)). If present, the branch is recorded as **concordant** for that gene; if absent, it is recorded as **discordant** (topology-induced absence under free-topology gene trees).

A key consequence of projection under missing taxa is **branch fusion**. For a given gene *g*, multiple distinct species-tree branches can yield identical projected splits, i.e., σ_g_(*b_1_*) = σ_g_(*b_2_*). Such branches form a **fusion group**, meaning they are indistinguishable on *T_g_.* Under fixed topology, SplitAligner can report these fused branches explicitly (e.g., as Bs1|Bs3|…) and aggregate branch-length information in a projection-aware manner.

Based on these definitions, SplitAligner produces three standardized outputs. First, it generates **fixed-topology** and **free-topology** branch-assignment/branch-length tables in a common species-branch coordinate system, while preserving branch-level missingness states (structural vs topology-induced). Second, it provides an optional **hybrid (mix-fixed) table** in which topology-induced missing entries in the free-topology table are filled by the corresponding fixed-topology values and explicitly flagged, enabling controlled quantification of discordance-driven missingness. Third, it outputs a **branch-wise concordance annotation** on the species tree, summarizing for each internal branch the fraction of decisive genes whose free-topology gene trees recover the projected split. These outputs together establish an infrastructure for defining, estimating, and comparing branch-specific quantities across loci under pervasive missing taxa and topological discordance.

### 2.2 Algorithm

SplitAligner: six-step split-alignment for gene–species branch mapping

#### Input

- Gene tree *G* with branch lengths (optional) and leaf set *L_G_*
- Species tree *S* with leaf set *L_S_*
- Optional parameters: maximum merge gap / mapping options (as used in implementation)

#### Output

- A gene-to-species **branch mapping matrix** *M*, where each gene-tree split is mapped to:
- a unique species-tree branch, or (ii) a fused species-branch set, or labeled as NA with a reason.

#### Step 1 — Harmonize taxa (missing-taxa trimming)

1. Let *L* = *L_G_* ∩*L_S_*.
2. Prune *G* and *S* to the shared leaf set *L*, producing *G’* and *S’*.
3. (Optional) If L is below a minimum threshold, mark this gene as insufficient and stop.

#### Step 2 — Extract gene-tree splits

4. From *G’*, enumerate internal branches and represent each branch as a **split** *gi* (a bipartition of L).
5. Record branch lengths for *g_i_* when available.
6. Store gene splits as an ordered list *G_splits_* = {*g_1_*,…*g_m_*}.

#### Step 3 — Extract species-tree splits

7. From S’, enumerate internal branches and represent each branch as a **split** *s_j_* (a bipartition of *L*).
8. Store species splits as an ordered list *S_splits_* = {*s_1_*,…*s_n_*}.

#### Step 4 — Split alignment (direct mapping)

9. For each gene split *g_i_* ∈ *G_splits_*:

- if there exists a species split *s_j_* ∈ *S_splits_* such that *g_i_* = *s_j_* (up to side ordering), then set 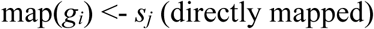
- Else, set map(*g_i_*) <- NA_candidate (unmapped at this step).

#### Step 5 — Fused-branch resolution (when direct mapping fails)

10. For each unmapped gene split *g_i_* (NA_candidate):

- Identify a set of **species splits** *F(g_i_)* ⊆ *S_splits_* whose union corresponds to the same clade boundary implied by *g_i_* on the pruned taxon set *L*.
- If such a set exists and satisfies the implementation’s validity rules, define 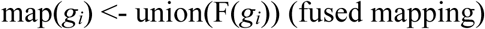
- Otherwise, keep *map(g_i_) <- NA* and assign an NA reason (Step 6). *(Implementation note: fused-branch length can be computed as the sum of constituent species-branch lengths when all are defined; if any constituent is NA, the fused length is NA.)*

#### Step 6 — Assign NA categories and export mapping matrix

11. For any *g_i_* still unmapped after Step 5, assign NA labels based on diagnostic rules, e.g.:

- **NA_topo:** topological incompatibility (no valid fused representation on *S’*)
- **NA_struct:** required species branches are structurally absent under the current taxon configuration / missingness
- **NA_fuse:** gene split corresponds to a fused branch where constituent parts are individually NA under the chosen bookkeeping scheme
12. Construct the mapping matrix *M* that records, for each *g_i_*:

- mapped species branch ID or fused-branch ID
- branch length(s) (if used)
- NA label (if applicable)
13. Output *M* for downstream counting and decomposition analyses.

## 3. Results

### 3.1 Projection under missing taxa

#### Fixed-topology mapping under taxon missingness reveals structural missingness and branch fusion

Under pervasive taxon missingness, direct comparison of branches across gene trees is ill-defined because each gene is inferred on a different taxon set. SplitAligner therefore defines branch identity on the species-tree backbone via splits and evaluates each branch *b* on a gene-specific taxon set *T_g_* using the projected split σ_g_(*b*) (Fig. 1A). If σ_g_(*b*) is degenerate, the branch is not evaluable for that gene, producing **structural missingness (NA_struct)**. Projection also makes explicit a second consequence of missing taxa: multiple species-tree branches can yield identical projected splits on *T_g_*, forming a **fusion group** that is indistinguishable for that gene (Fig. 1B). Under fixed-topology (species-tree–constrained) gene trees, SplitAligner reports the corresponding **fused branch identity** (e.g., Bs1|Bs3) and aggregates branch-length information in a projection-aware manner, while marking the original member branches as **NA_fuse** (Fig. 1B-C). This establishes a consistent branch coordinate system for downstream branch-wise analyses even when terminal taxa are missing.

**Figure 1.**
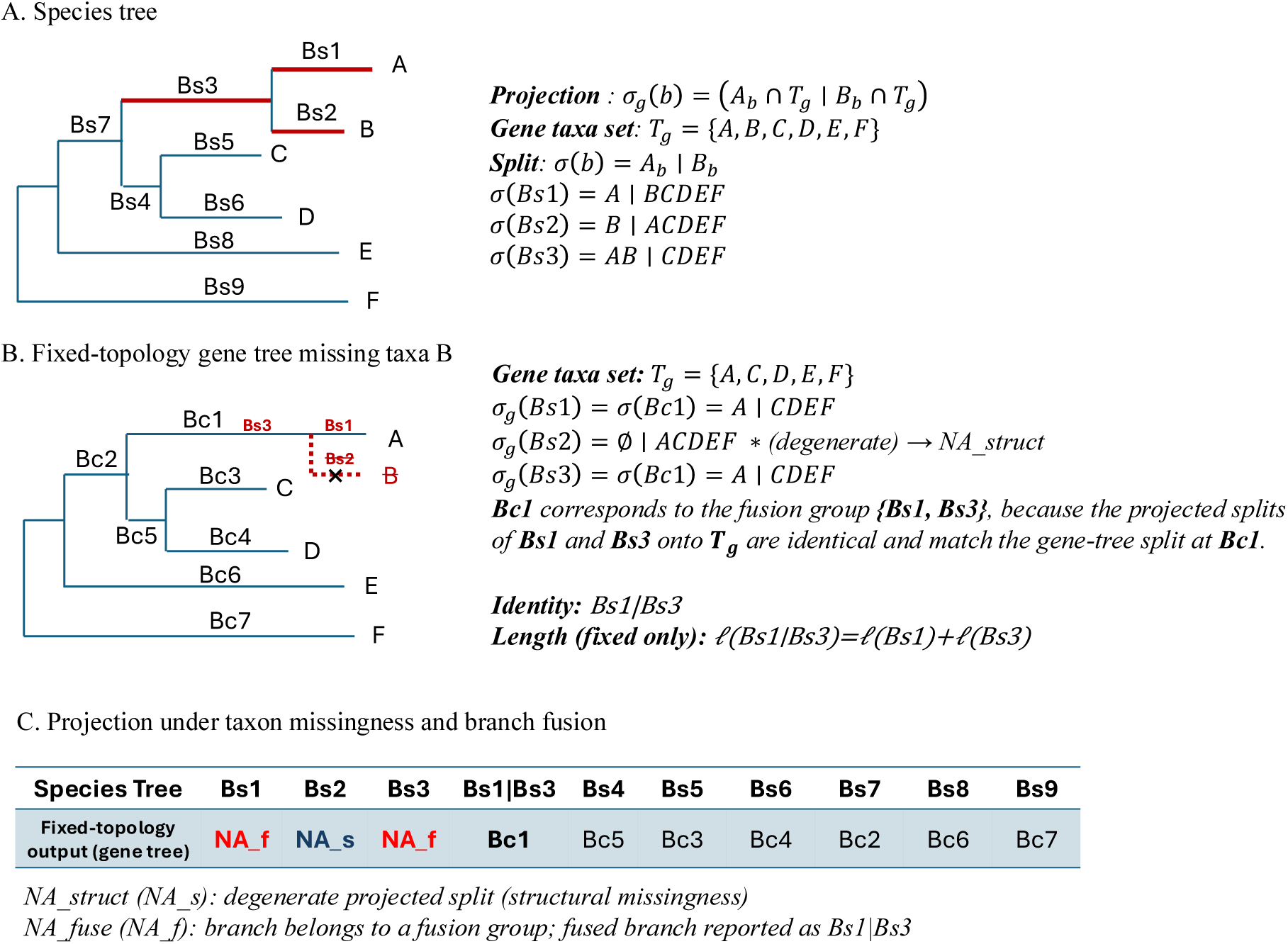
Projected splits define branch identity under taxon missingness. Projecting species-tree splits onto the gene-specific taxon set can render some branches structurally missing (degenerate projected splits) and can collapse multiple species-tree branches into an indistinguishable fusion group, represented by a composite branch identity (e.g., **Bs1|Bs3**) under fixed topology. Branches belonging to a fusion group are marked as **NA_fuse** at their original species-branch rows, while the composite identity is reported as an additional branch (e.g., ***Bs1|Bs3***).

### 3.2 Discordance adds topology-induced missing

#### Free-topology mapping separates topology-induced missingness from structural missingness

Taxon missingness alone does not explain all missing entries in branch-assignment tables when gene trees are inferred with free topology. For a given species-tree branch *b*, if the projected split σ_g_(*b*) is non-degenerate, gene *g* is **decisive** for *b*; in this case, absence of σ_g_(*b*) from the free-topology gene tree indicates **topology-induced missingness (NA_topo)** driven by discordance rather than lack of evaluability (Fig. 2A-B). SplitAligner distinguishes this case from **structural missingness (*NA_struct*)**, where σ_g_(*b*) is degenerate due to insufficient taxon coverage. This separation is critical because NA_topo is expected to be concentrated on internally discordant regions of the species tree, whereas NA_struct primarily reflects gene-specific taxon coverage (Fig. 2B-C). By retaining these missingness states explicitly in the free-topology table, SplitAligner enables quantitative analyses of discordance-induced bias without conflating it with coverage limitations.

**Figure 2.**
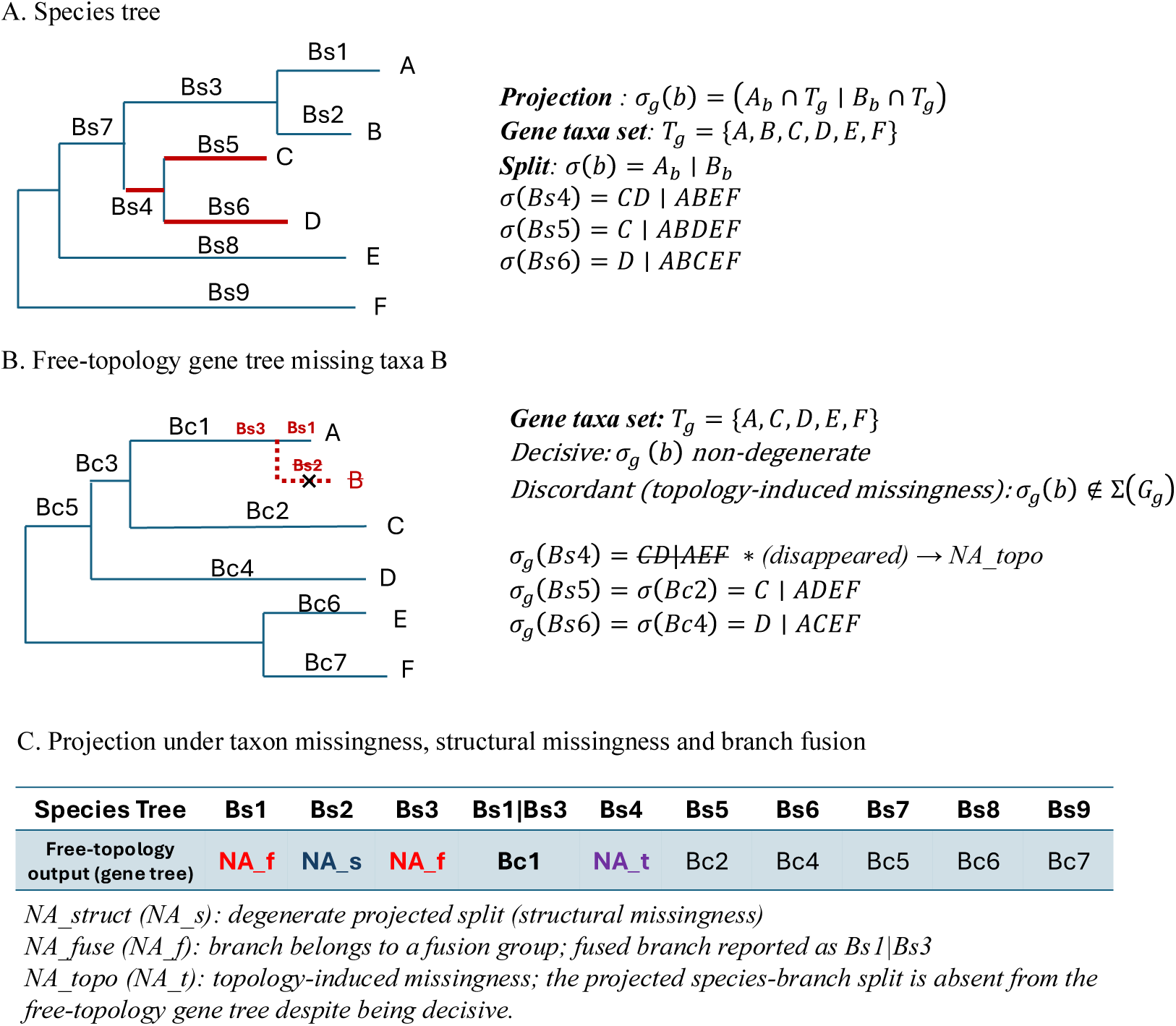
Topology-induced missingness under free-topology gene trees. Under taxon missingness, projected species-tree splits remain the reference for branch identity; a branch is **decisive** for a gene if its projected split is non-degenerate. Under **free-topology** gene trees, a decisive projected split may nevertheless be absent from the gene tree, yielding **NA_topo** (topology-induced missingness). In contrast, degenerate projected splits produce **NA_struct** (structural missingness), and branches belonging to a fusion group are marked as **NA_fuse** with the fused-branch identity represented by an additional composite branch (e.g., Bs1|Bs3). *Gene trees may also contain additional splits that have no counterpart in the species tree; SplitAligner focuses on branch-wise evaluation of projected species-tree splits in a common backbone coordinate system*.

### 3.3 Branch-wise concordance score and its distribution on internal nodes of a real dataset

To summarize discordance in a branch-resolved manner on the species-tree backbone, we computed a branch-wise concordance score (*Support*) for each internal branch of the 302-mammal species tree. For a given branch *b*, SplitAligner evaluates, gene by gene, whether the projected species-tree split of *b* is recovered in the corresponding free-topology gene tree.

Because taxon coverage varies across genes, we normalize concordance by the number of genes for which branch *b* is evaluable under the fixed-topology mapping, yielding *Support(b)* = *100* × *n_unfix_ (b)* / *n_dec_(b)*. Support is conceptually distinct from bootstrap values or posterior probabilities: it directly quantifies the empirical frequency with which gene trees recover each species-tree branch, conditional on being comparable under missing taxa. We visualized Support on a compact hominoid subtree (Fig. 3A) to provide an externally validated sanity check. At the well-known human–chimpanzee–gorilla divergence, SplitAligner estimates support = 73% for the human+chimpanzee branch, consistent with extensive incomplete lineage sorting at this short internode (Fig. 3B). Extending this annotation genome-wide, Support varies substantially across the mammal phylogeny (Fig. 3C), indicating that discordance-driven absence is not uniformly distributed among internal branches. This branch-wise concordance annotation provides a practical diagnostic for identifying internally discordant regions of the species tree and motivates downstream analyses of topology-induced missingness and bias.

**Figure 3.**
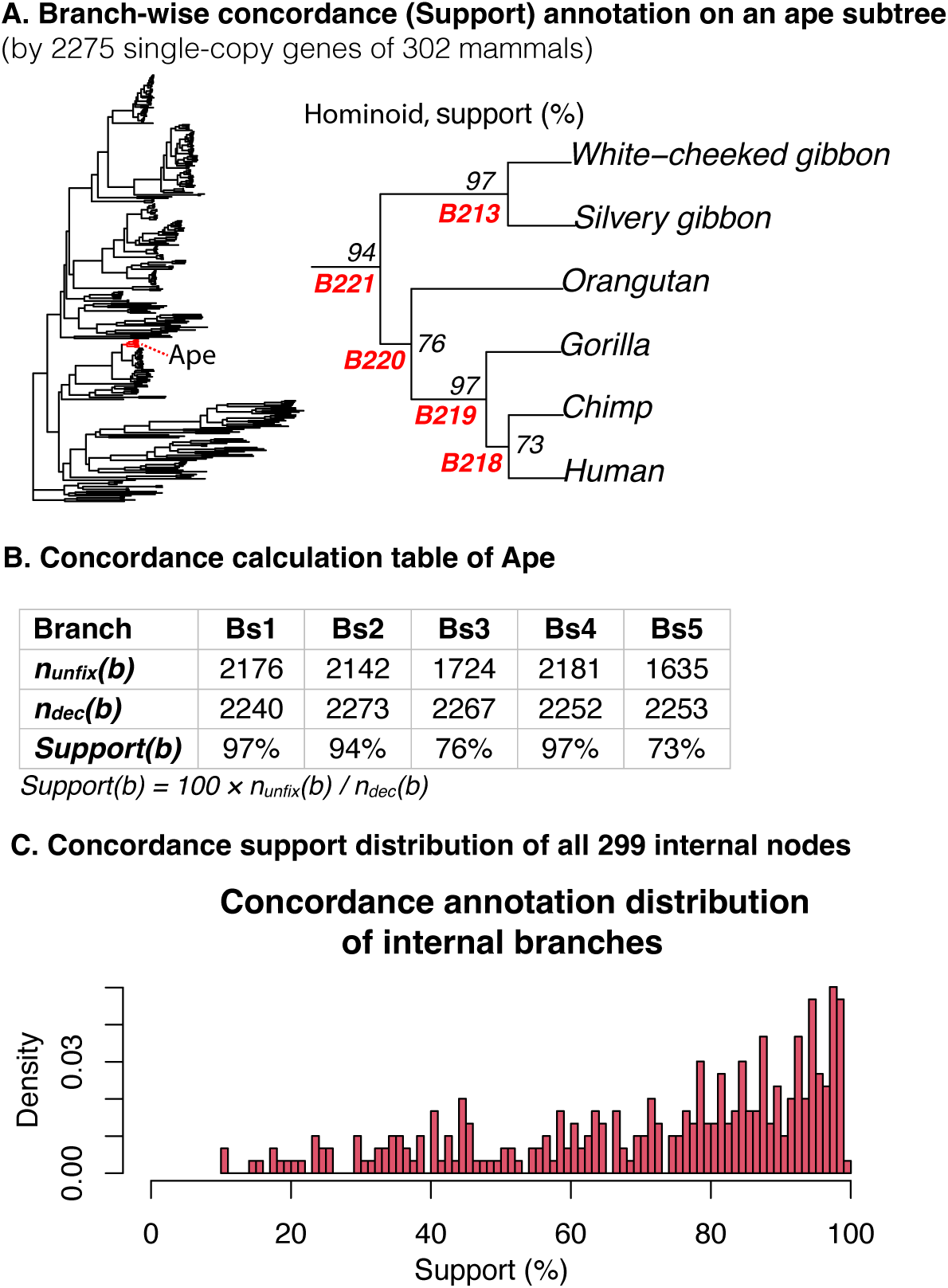
Branch-wise concordance (Support) annotation on the species-tree backbone. (A) SplitAligner annotates each internal species-tree branch with a branch-wise concordance score, Support (%), computed from 2,275 single-copy genes in the 302-mammal dataset. A compact hominoid subtree highlights the classic human–chimpanzee–gorilla internode, where the human+chimpanzee branch shows Support = 73%, consistent with extensive incomplete lineage sorting at this divergence. (B) For a given species-tree branch *b*, Support is defined as the fraction of **decisive** genes whose free-topology gene trees contain the **projected split** of *b*; a gene is decisive for *b* if the projected split σ_g_(*b*) is non-degenerate on the gene-specific taxon set. (C) Genome-wide distribution of Support across all internal branches of the mammal species tree. Support is conceptually distinct from bootstrap or posterior probabilities because it summarizes empirical gene-tree concordance with each species-tree branch under explicit control of missing taxa.

Building on the branch-wise concordance annotation (Fig. 3), we quantified how gene-tree/species-tree discordance translates into systematic missingness under free-topology mapping. For each species-tree branch b, we distinguished structural missingness (degenerate projected splits) from topology-induced missingness, where a branch is decisive but its projected split is absent from the free-topology gene tree. Across internal branches, topology-induced missingness increases strongly with branch discordance (Fig. 4A), demonstrating that discordance-driven absence is concentrated on specific internodes rather than uniformly distributed. Stratifying branches by concordance further reveals a marked separation in topology-induced missingness (Fig. 4B), supporting the view that free-topology mapping yields branch-dependent evaluability and can induce systematic bias in downstream branch-wise summaries if discordance is ignored. The most discordant internodes (Fig. 4C) provide concrete targets for diagnosing and interpreting discordance hotspots in large phylogenomic datasets.

**Figure 4.**
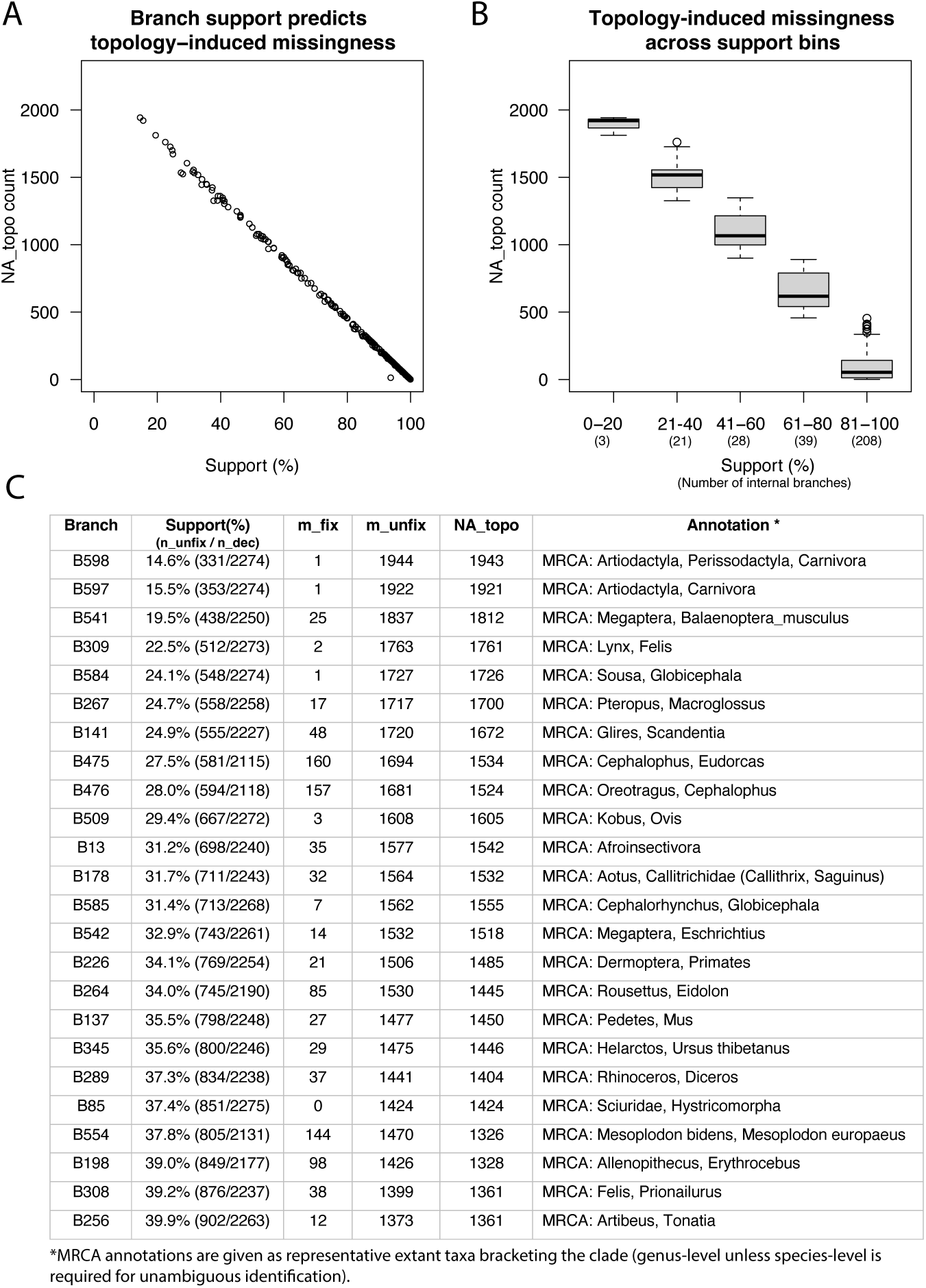
Branch support predicts topology-induced missingness across internal branches. (A) Each point represents one internal branch of the 302-species mammal species tree. The x-axis shows Support (%), defined as *100* × *n_unfix_* / *n_dec_*, where *n_dec* is the number of decisive genes under the fixed-topology baseline and *n_unfix* is the number of those genes that recover the projected split under free-topology mapping. The y-axis shows *NA_topo* count, defined as the excess missing mappings under free-topology relative to fixed-topology (*m_unfix_* − *m_fix_*). (B) Boxplots of *NA_topo* across support bins (0–20, 21–40, 41–60, 61–80, 81–100%), with the number of internal branches in each bin shown in parentheses. (C) Branches with the largest *NA_topo*. For each branch we report Support (%; *n_unfix_* / *n_dec_*), *m_fix_*, *m_unfix_*, NA_topo (*m_unfix_* − *m_fix_*), and a representative clade label (MRCA of listed taxa).

### 3.4 Decomposing missingness into structural, fusion, and topology-induced components

Building on the branch-wise concordance patterns (Figs. 3–4), we next decomposed missing mapping outcomes into three mechanistic components: **structural missingness** (NA_struct; degenerate projected splits driven by taxon coverage), **fusion-row missingness** (NA_fuse; indistinguishable adjacent species-tree branches under a gene-specific taxon set, with signal reassigned to composite fused branches), and **topology-induced missingness** (NA_topo; decisive projected splits that are absent from free-topology gene trees). This decomposition provides a concrete accounting of why a species-tree branch can be “missing” under gene-tree discordance and missing taxa, and clarifies how these missingness modes differ between fixed-topology and free-topology mapping.

Across **internal branches**, NA_topo emerged as the dominant difference between fixed and free mapping: NA_topo was absent by definition under the fixed-topology baseline but became abundant under free-topology mapping, whereas NA_struct remained invariant and NA_fuse changed only modestly (Fig. 5A). In contrast, **terminal branches** showed no topology-induced excess missingness, with missingness almost entirely attributable to NA_struct and NA_fuse (Fig. 5B). Consistent with these patterns, internal and terminal counts sum exactly to the genome-wide totals (Fig. 5C), confirming that discordance redistributes mapping outcomes rather than introducing ambiguous bookkeeping. Finally, focusing on all **low-support internodes** (Support < 40%; n=24) highlights that their missingness is overwhelmingly composed of *NA_topo* under free-topology mapping (Fig. 5D), providing a direct mechanistic explanation for the strong support–missingness relationship observed in Fig. 4.

**Figure 5.**
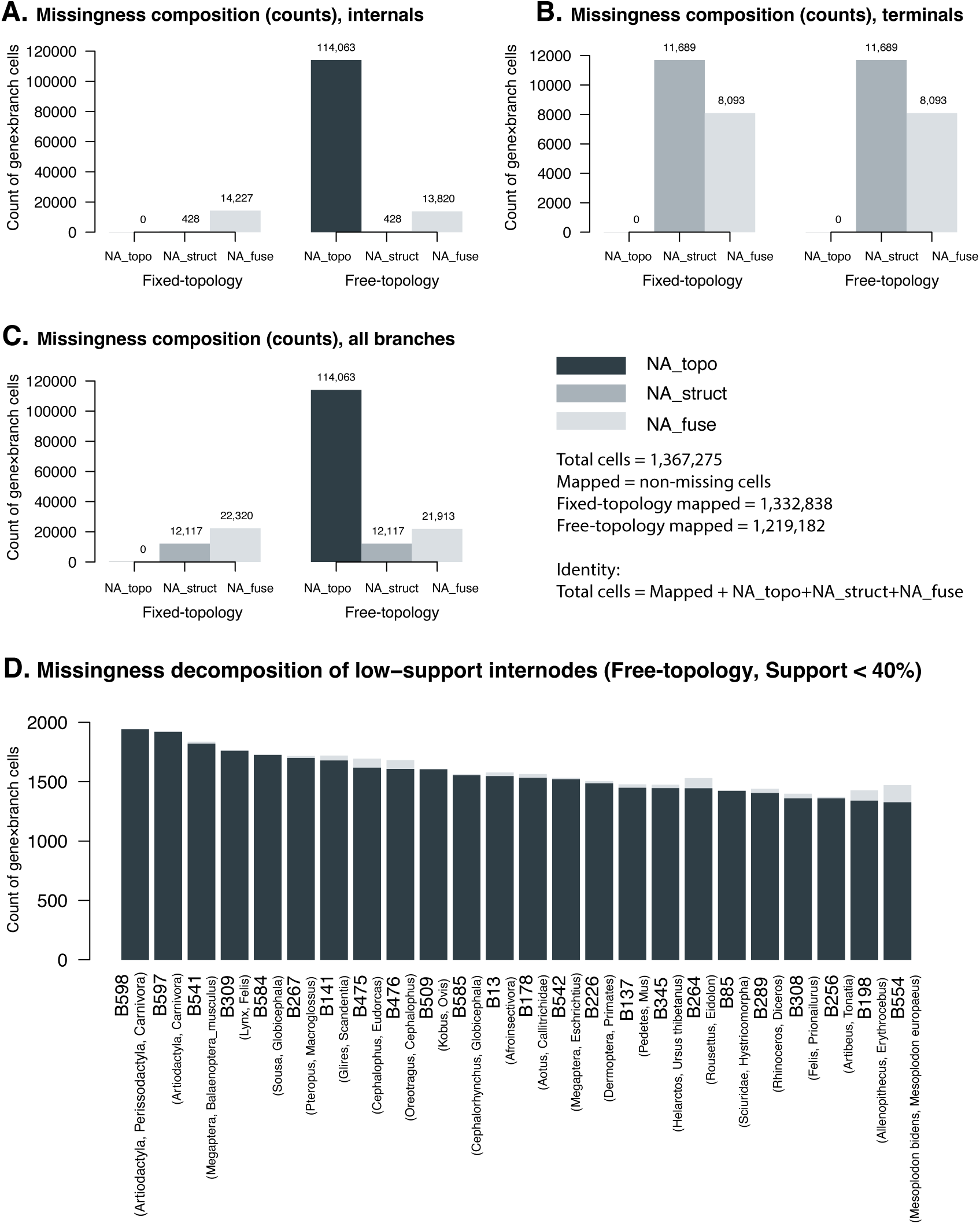
Decomposing missingness into topology-induced, structural, and fusion components. (A–C) Missingness composition (counts of gene×branch cells) under fixed-topology and free-topology mapping, shown separately for internal branches (A), terminal branches (B), and all branches (C). Each cell is classified as Mapped (non-missing), NA_struct (degenerate projected split due to taxon coverage), NA_fuse (fusion-row missingness at original species-branch rows; corresponding signal may be represented on composite fused-branch rows), or NA_topo (decisive projected split absent from the free-topology gene tree). Internal and terminal counts sum exactly to the all-branch totals; the matrix-level accounting identity is shown in-panel. (D) Missingness decomposition for low-support internodes under free-topology mapping (Support < 40%; n = 24). Each bar represents one internode, partitioning gene×branch cell counts into NA_topo, NA_struct, and NA_fuse. Parenthetical labels denote representative clades (MRCA of listed taxa).

Importantly, the three missingness types form an explicit bookkeeping system. For each mapping cell (gene *g* × species-tree branch *b*) under a fixed taxon set, projected-split evaluation yields one of four mutually exclusive outcomes: **Mapped**, **NA_struct**, **NA_fuse**, or **NA_topo**. Thus, at the matrix level we have the accounting identity

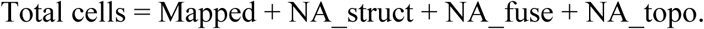

Moreover, under free-topology mapping, topology-induced missingness reflects a one-to-one replacement of projected species splits by alternative gene-tree splits: for each gene *g*, the number of **unmapped gene-tree splits** equals the number of **NA_topo** projected splits (i.e., decisive projected splits absent from the gene tree). This provides an internal consistency check that discordance redistributes splits rather than removing signal.

## 4. Method

### 4.1 Genome alignments and quality filtering

We downloaded mammalian coding-gene alignments generated by the Hiller Lab (TOGA-based orthology inference on the human hg38 reference assembly) (*9*). The resource contains a catalog of 17,434 coding-gene alignments from 517 placental mammal genome assemblies representing 442 species. To define a consistent species-level taxon set, we retained species with fewer than 1,800 missing genes across the catalog. When multiple assemblies (e.g., subspecies) were available for the same species, we kept a single representative assembly with the fewest missing genes. This procedure resulted in a 302-species placental mammal dataset used in this study.

We further defined a high-confidence single-copy subset for species-tree inference and SplitAligner evaluation, following Wu et al. (2017) (*10*). Specifically, we retained genes with alignment length > 2,000 bp and missing sequences in fewer than 30 of the 302 species, resulting in 2,275 genes, which are released with this study.

### 4.2 Species tree inference

A reference species tree was inferred from the 2,275 single-copy genes. For each gene, we inferred an unconstrained (free-topology) gene tree using IQ-TREE2 with automatic substitution-model selection and partitioning by codon position (1st+2nd versus 3rd positions) (*11*). The species tree was then estimated using ASTRAL under the multispecies coalescent model from the collection of inferred gene trees (*12*).

### 4.3 Fixed-topology branch-length estimation and SplitAligner inputs

SplitAligner takes gene trees and a reference species tree as input and supports both fixed-topology and free-topology mapping. For **fixed-topology** mapping, we constrained each gene tree topology to the reference species tree and estimated **gene-wise branch lengths on the fixed backbone** using BaseML (PAML) (*4*). Unless otherwise stated, branch lengths were estimated under the GTR+Γ+I model (*13–15*). This produces a fixed-topology branch-length table in which branch identity is aligned to the species-tree coordinate system by construction, while allowing gene-specific missing taxa to induce degenerate projections (NA_struct) and fusion groups (NA_fuse).

For **free-topology** mapping, SplitAligner maps unconstrained gene trees (IQ-TREE2) onto the same species-tree branch coordinate system via split-based branch mapping. By comparing fixed-topology and free-topology mapping outcomes, SplitAligner separates missingness due to taxon coverage (NA_struct and NA_fuse) from missingness induced by topological discordance (NA_topo), as quantified in Figs. 4–5.

### 4.4 SplitAligner mapping and output tables

SplitAligner maps gene-tree information onto a fixed species-tree branch coordinate system using split-based branch identities. For a reference species tree and a gene tree with gene-specific taxon set *T_g_*, each species-tree internal branch induces a species split, which

SplitAligner **projects** onto *T_g_* by restricting both sides of the split to taxa present in the gene. If the projected split becomes trivial due to insufficient taxon coverage (e.g., one side is empty), the branch is not evaluable for that gene and is labeled **NA_struct** (structural missingness).

Under fixed-topology gene trees (species-tree–constrained), projected splits naturally reveal **branch fusion** induced by taxon missingness: multiple adjacent species-tree branches may yield identical projected splits on the same *T_g_*. SplitAligner represents such cases by reporting an explicit **composite fused-branch identity** (e.g., B1|B3|…) and marking the corresponding original branch rows as NA_fuse (fusion-row missingness). When branch lengths are used under fixed topology, fused-branch lengths are aggregated from the constituent species-tree branches.

Under free-topology gene trees, a branch can be evaluable (non-degenerate projection) yet its projected species split may be absent from the gene tree due to discordance. SplitAligner labels this outcome as NA_topo (topology-induced missingness), separating discordance-driven absence from NA_struct driven by taxon coverage alone. SplitAligner outputs fixed-topology and free-topology gene×branch mapping tables (TSV), optionally including composite fused-branch columns, together with NA_struct / NA_fuse / NA_topo labels for downstream counting and decomposition analyses.

### 4.5 Branch-wise concordance (Support) and accounting identities

We quantified branch-wise gene-tree concordance on the species-tree backbone using a **Support** score for each internal species-tree branch *b*. Branches labeled as NA_fuse are excluded from **decisiveness** at the member-branch level because missing taxa can collapse multiple adjacent species-tree branches into an indistinguishable fusion group on *T_g_*, preventing unique attribution to any single constituent branch. For each gene *g*, SplitAligner evaluates whether branch *b* is decisive under the gene-specific taxon set *T_g_*, i.e., whether the projected species-tree split of *b* on *T_g_* is non-degenerate and yields a unique species-branch identity under the fixed-topology baseline (i.e., not labeled as NA_struct or NA_fuse). Among decisive genes, a gene is considered supporting branch *b* if SplitAligner yields a non-missing mapping result on *b* in the free-topology table (equivalently, the projected split of *b* is present in the free-topology gene tree).

Let *N* be the number of genes. For branch *b*, define *m_fix_(b)* and *m_unfix_(b)* as the numbers of missing mapping outcomes for *b* in the fixed-topology and free-topology tables, respectively.

The number of decisive genes for branch *b* is

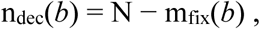

and the number of genes that recover the projected split of *b* under free-topology mapping is

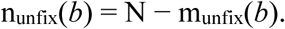

Here, *n_unfix_(b)* denotes the number of genes whose free-topology gene trees yield a non-missing mapping on branch *b* (i.e., recover the projected split of *b*). We then define branch-wise concordance (Support) as

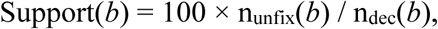

and branch discordance as

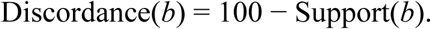

In this formulation, Support is a cross-gene concordance measure on a fixed branch coordinate system, conditional on decisiveness under taxon coverage, and is therefore distinct from bootstrap support or ASTRAL-derived branch support measures.

SplitAligner’s NA labeling yields an explicit bookkeeping system at the gene×branch matrix level. Each matrix cell belongs to exactly one of four mutually exclusive states: Mapped, NA_struct, NA_fuse, or NA_topo. Accordingly, the total number of gene×branch cells satisfies the accounting identity

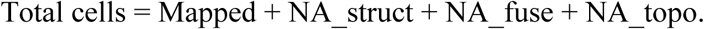

In addition, under free-topology mapping, topology-induced missingness corresponds to a one-to-one replacement of projected species splits by alternative gene-tree splits: for each gene under a fixed taxon set, the number of **unmapped gene-tree splits** equals the number of NA_topo projected splits. This relationship provides an internal consistency check that discordance redistributes split support rather than removing signal.

Optionally, we provide a diagnostic hybrid table in which entries missing under free-topology mapping (NA_topo) are imputed using the corresponding fixed-topology estimates and explicitly flagged. This hybrid output is intended for controlled comparisons (e.g., *Fix* vs *Unfix bias quantification*) rather than as a replacement for free-topology mapping.

## 5. Implementation and availability

SplitAligner is implemented as a lightweight, file-based command-line pipeline. The core inputs are a reference species tree (*Newick*) and a set of per-gene gene trees (*Newick*; one tree per locus), and the core outputs are branch-mapping matrices in tab-delimited format for downstream analyses.

### Core workflow

For each gene, SplitAligner (i) extracts splits from the gene tree and the species tree, (ii) projects species-tree splits onto the gene-specific taxon set, (iii) performs split matching to map gene-tree splits onto the species-tree branch coordinate system, and (iv) reports mapped branches, composite fused branches (when applicable), or missing outcomes. The pipeline produces two primary branch-mapping tables: a **fixed-topology table** (branch lengths estimated on a fixed backbone) and a **free-topology table** (unconstrained gene trees), which together enable separation of taxon-coverage-driven missingness (NA_struct / NA_fuse) from discordance-driven missingness (NA_topo).

### Outputs

SplitAligner writes (1) gene×branch matrices for fixed-topology and free-topology mapping, (2) optional composite fused-branch columns (e.g., B1|B3) for explicit representation of fusion groups, and (3) optional species-tree annotations (e.g., branch-wise Support). Output tables are plain-text TSV files designed for easy counting and visualization.

### Optional post-processing utilities (for Fig. 5 decomposition)

We provide two small utilities that operate on the SplitAligner output matrices without altering the underlying split mapping: Extract_NA_fuse.pl labels fusion-row missing entries (NA_fuse) based on the presence of composite fused-branch columns, and Confirm_NA_structure.pl labels structural missingness (*NA_struct*) and topology-induced missingness (NA_topo) by cross-referencing fixed- and free-topology matrices under identical branch headers. These utilities are optional and are used here to generate the missingness decomposition shown in Fig. 5.

### Computational complexity

Given precomputed splits, per-gene mapping scales approximately linearly with the number of genes and the number of splits per gene; fusion resolution is evaluated only when direct matching fails. The implementation is suitable for large phylogenomic datasets with thousands of loci.

### Availability

Source code and documentation are available at https://github.com/wujiaqi06/SplitAligner.

## 6. Discussion

**SplitAligner addresses a practical but often implicit problem in phylogenomics: branch identity becomes unstable when gene trees and species trees differ in both topology and taxon sampling.** By formulating branch mapping in a split-based coordinate system, SplitAligner makes branch comparability explicit under gene-specific taxon sets. In particular, the notion of **projected species-tree splits** provides a deterministic basis for evaluating whether a gene tree can be meaningfully compared to a given species-tree internode, even when many taxa are missing. This resolves a common ambiguity in downstream branch-wise analyses, where “missing” values are frequently treated as a single category despite arising from distinct mechanisms.

A central outcome of this framework is the explicit **decomposition of missingness** into NA_struct, NA_fuse, and NA_topo, together with a bookkeeping identity at the gene×branch matrix level. NA_struct captures degeneracy of projected splits driven by taxon coverage, NA_fuse captures cases where multiple adjacent species-tree branches become indistinguishable after projection and are represented by composite fused branches, and NA_topo captures discordance-driven absence of decisive projected splits on free-topology gene trees. This decomposition clarifies why free-topology mapping can introduce systematic, branch-specific missingness beyond taxon coverage alone, and it provides internal consistency checks that help distinguish discordance effects from incomplete sampling or implementation artifacts.

SplitAligner’s **Support** annotation complements, rather than replaces, traditional measures of phylogenetic confidence such as bootstrap support or ASTRAL-derived local measures.

Bootstrap quantifies stability under resampling within a single-tree model, whereas Support summarizes **cross-gene concordance on a fixed species-tree branch coordinate system**, conditional on decisiveness under taxon coverage. In this sense, Support is directly aligned with the needs of branch-wise comparative analyses across loci: it highlights internodes that are repeatedly recovered as projected splits across free-topology gene trees and anticipates where discordance is expected to manifest as topology-induced missingness (NA_topo). **Recommended usage** is therefore to interpret Support jointly with the missingness decomposition—especially on internal branches—rather than treating all missing mappings as equivalent. SplitAligner is related to split-based summaries such as PhyParts and concordance-factor approaches (e.g., gCF/sCF), but differs in explicitly defining branch identity under gene-specific taxon sets and decomposing missingness into NA_struct/NA_fuse/NA_topo under a matrix-level accounting identity (*6–8*). In particular, SplitAligner interprets missing-taxa–driven ambiguous multi-mapping as branch fusion with explicit composite identities, enabling standardized gene×branch tables for downstream analyses rather than node-centric concordance summaries.

Notably, the lowest-support internodes identified by SplitAligner correspond to several well-known “difficult” nodes in mammalian phylogeny that have been repeatedly discussed in the literature, including deep placental splits and rapid radiations within Laurasiatheria and Cetartiodactyla, as well as lineage-rich clades such as primates, bats, and odontocetes. These internodes are not merely low-confidence in a generic sense: SplitAligner shows that their apparent instability is accompanied by elevated NA_topo under free-topology mapping, indicating that discordance manifests as topology-driven absence of projected species splits rather than as simple taxon-coverage loss (NA_struct) or fusion-row bookkeeping (NA_fuse). In this way, SplitAligner provides a mechanistic, branch-resolved decomposition of where and how discordance erodes branch comparability across loci, offering a practical diagnostic layer complementary to bootstrap or coalescent-based support values. We emphasize that SplitAligner does not re-infer phylogenetic relationships for these nodes; rather, it provides an infrastructure for making branch identity and branch-wise discordance/missingness explicit under heterogeneous taxon sampling.

Finally, we note practical considerations and extensions:

- **Scope and assumptions.** SplitAligner assumes a fixed reference species tree and gene trees inferred on (possibly different) taxon subsets. Mapping is performed at the level of splits and is intended as a branch-coordinate infrastructure rather than a network/reticulation inference method.
- **Interpretation of fusion.** Fusion is reported as a branch-row bookkeeping construct: NA_fuse indicates indistinguishability of adjacent species-tree branches under projection, with signal potentially reassigned to composite fused branches. Users should interpret NA_fuse accordingly when summarizing branch-level signals.
- **Extensions.** The projected-split coordinate system and accounting framework can be extended to alternative reconciliation rules, additional diagnostics, or explicit network representations, while preserving the key goal of comparable branch identity under missing taxa and discordance.

Future work may leverage the distribution of unmapped gene-tree splits (complements of NA_topo under fixed taxon sets) to characterize alternative split support around highly discordant internodes, including classical three-taxon conflicts.

## Data availability

All data products generated in this study—including the 2,275 single-copy gene trees, gene×branch mapping tables with NA_struct/NA_fuse/NA_topo labels, branch-wise Support summary tables (including the full table underlying Fig. 4C), and the annotated species tree—are available at the SplitAligner GitHub repository (https://github.com/wujiaqi06/SplitAligner) in standard plain-text formats (e.g., *Newick*, *TSV*), together with scripts to reproduce the figures and key analyses.

## Author Contributions

J.W. conceived the study, developed the SplitAligner algorithm, implemented the software, performed all computational analyses, generated the figures, and wrote the manuscript.

## Acknowledgement

This study was supported by Grant-in-Aid for Scientific Research (B) 23K23950 from the Japan Society for the Promotion of Science. I thank ChatGPT (OpenAI) for assistance with language editing, manuscript organization, and figure-caption polishing. All scientific decisions, analyses, and interpretations were made by the author.

## Definitions

Species tree (*S*): a fixed reference phylogeny representing species relationships, used as the target structure for split mapping.
Gene tree (*G_g_*): a phylogenetic tree inferred from a single gene or locus *g*, potentially with missing taxa and topological discordance relative to *S*.
Fixed-topology gene tree (species-tree–constrained gene tree): a gene tree whose topology is constrained to the species tree; only branch lengths are estimated on this fixed backbone.
Free-topology gene tree (unconstrained gene tree): a gene tree inferred without topological constraints.
Gene-specific taxon set (*T_g_*): the set of taxa present in gene *g*.
Split (bipartition): a bipartition of taxa induced by an internal branch of a tree. Species-tree split (σ(*b*)) the split induced by an internal branch *b* of the species tree *S*. Gene-tree split the split induced by an internal branch of a gene tree *G_g_*.
Split set (Σ(*G_g_*)): the set of all gene-tree splits (bipartitions) induced by the internal branches of *G_g_*.
Projected split (σ_g_(*b*)): the restriction (projection) of a species-tree split σ(*b*) onto the gene-specific taxon set *T_g_*, yielding σ_g_(*b*) = (*A_b_* ∩ *T_g_* | *B_b_* ∩ *T_g_*).
Degenerate projected split: a projected split that collapses to a trivial partition (e.g., one side becomes empty) due to insufficient taxon coverage.
Taxon missingness: missing taxa in a gene tree relative to the species tree.
Gene×branch cell: one entry in a gene × branch mapping table for gene *g* and species-tree branch *b*.
Structural missingness (NA_struct): the absence of informative signal for a species-tree branch *b* in gene *g* because the projected split σ_g_(*b*) is degenerate (the branch is not evaluable for that gene).
Branch fusion (fusion group): under a given taxon set *T_g_*, multiple distinct species-tree branches can yield identical projected splits, i.e., σ_g_(*b_1_*) = σ_g_(*b_2_*), making these branches indistinguishable at the member-branch level.
Composite fused-branch identity (b_1_|b_2_|…): an explicit branch label representing a fusion group under a given *T_g_*, used as an additional composite row/column to store the fused signal when member branches are indistinguishable.
Fusion-row missingness (NA_fuse): the missing label assigned to member-branch rows when *b* belongs to a fusion group under T_g; the corresponding signal may be represented on an additional composite fused-branch row (e.g., b_1_|b_2_|…).
Decisive (gene for a species-tree branch): a gene *g* is decisive for branch *b* if σ_g_(*b*) is non-degenerate and yields a unique species-branch identity under the fixed-topology baseline (i.e., not labeled as NA_struct or NA_fuse).
Concordant split: a decisive projected species-tree split σ_g_(*b*) that is present in the corresponding gene tree (i.e., σ_g_(*b*) ∈ Σ(*G_g_*)).
Discordant split: a decisive projected species-tree split σ_g_(*b*) that is absent from the corresponding gene tree; its absence implies support for alternative splits in that region (often incompatible with σ_g_(*b*)).
Concordant (gene) with branch b: the gene tree contains the projected split of branch *b* (i.e., σ_g_(*b*) is concordant for gene *g*).
Discordant (gene) with branch b: the gene tree does not contain the projected split of branch *b* despite gene *g* being decisive for *b* (i.e., σ_g_(*b*) is discordant for gene *g*).
Topology-induced missingness (NA_topo): for a decisive gene–branch pair, the mapping is missing under free-topology inference because σ_g_(*b*) is absent from the free-topology gene tree (discordance-driven absence rather than coverage-driven non-evaluability).

